# Somatic junctions connect microglia and developing neurons

**DOI:** 10.1101/2021.03.25.436920

**Authors:** Csaba Cserép, Anett D. Schwarcz, Balázs Pósfai, Zsófia I. László, Anna Kellermayer, Miklós Nyerges, Zsolt Lele, István Katona, Ádám Dénes

**Author notes:** Lead Contact and Correspondence to: Ádám Dénes.

## Abstract

Microglia are the resident immune cells of the brain with multiple homeostatic and regulatory roles. Emerging evidence also highlights the fundamental transformative role of microglia in brain development. While tightly controlled, bi-directional communication between microglia and neuronal progenitors or immature neurons has been postulated, the main sites of interaction and the underlying mechanisms remain elusive. By using correlated light and electron microscopy together with super-resolution imaging, here we provide evidence that microglial processes form specialized nanoscale contacts with the cell bodies of developing and immature neurons throughout embryonic, early postnatal and adult neurogenesis. These early developmental contacts are highly reminiscent to somatic purinergic junctions that are instrumental for microglia-neuron communication in the adult brain. We propose that early developmental formation of somatic purinergic junctions represents an ideal interface for microglia to monitor the status of developing neurons and to direct prenatal, early postnatal and adult neurogenesis.

## Introduction

In the last couple of years, a series of breakthrough discoveries demonstrated that microglia, the brain’s resident immune cells are not only central players in various pathophysiological processes underlying brain disorders (Salter and Stevens, 2017), but also have essential physiological functions in the healthy brain (Kierdorf and Prinz, 2017; Li and Barres, 2018). Microglial progenitors originate from the yolk sac and appear in the CNS as early as the 5th gestational week in humans and embryonic day 9 (E9) in mice (De et al., 2018; Ginhoux et al., 2010; Verney et al., 2010). Accordingly, compelling evidence substantiates the critical importance of microglia in brain development (Lenz and Nelson, 2018; Mosser et al., 2017; Thion and Garel, 2017; Thion et al., 2018). The complexity of intercellular interactions in the central nervous system (CNS) emerges from a precisely regulated developmental program, which involves neuro- and gliogenesis, migration of postmitotic neuroblasts, formation and refinement of synapses and other forms of cell-cell connections as well as brain vasculature maturation (Martynoga et al., 2012; Silbereis et al., 2016; Urban and Guillemot, 2014). In accordance with the complexity of brain development, multiple studies confirmed that microglia regulate both embryonic and adult neurogenesis (Cunningham et al., 2013; Diaz-Aparicio et al., 2020; De Lucia et al., 2016; Sato, 2015; Sellner et al., 2016; Sierra et al., 2010; Walton et al., 2006), direct neuronal differentiation and migration (Aarum et al., 2003a), contribute to synaptogenesis and refinement (Gunner et al., 2019; Paolicelli et al., 2011; Rodríguez-Iglesias et al., 2019; Schafer et al., 2012) and drive the formation of cortical layers (Ueno et al., 2013). Microglia are also responsible for the removal of apoptotic neurons (Marín-Teva et al., 2004; Sierra et al., 2010), activity-dependent synapse formation (Parkhurst et al., 2013), synapse and spine remodeling (Weinhard et al., 2018) and axonal guidance (Pont-Lezica et al., 2014; Squarzoni et al., 2014). Although all of these developmental processes necessitate spatiotemporally coordinated interactions between microglia, neuronal progenitors and immature neurons, our understanding about the key sites of intercellular communication and underlying mechanisms in the developing brain has remained limited.

Communication between neurons and microglia takes place via multiple routes including indirect intercellular interactions through soluble messengers and direct, anatomically defined membrane-to-membrane contacts (Cserép et al., 2021). For example, microglia have been shown to modulate both developmental and adult neurogenesis through the extracellular release of cytokines (IL-1β, IL-6, TNF-α, IFN-γ, or TGF-β; Battista et al., 2006; Butovsky et al., 2006; Shigemoto-Mogami et al., 2014) and growth factors (Araki et al., 2020). In line with this, specialized, direct interactions such as those formed between microglia and neuronal synapses are required to shape brain circuit formation. For example, by contacting the dendrites of developing neurons with preference for more distal neurites (Chugh and Ekdahl, 2016), microglia induce spine outgrowth and synaptogenesis (Miyamoto et al., 2016). During synaptic pruning, microglial processes engulf and phagocytose non-essential synapses, which is also indispensable for proper activity-dependent brain circuit maturation (Paolicelli et al., 2011; Schafer et al., 2012; Weinhard et al., 2018). Development of axon initial segments and the inhibition of axonal outgrowth also take place via direct microglial contacts on growth cones (Baalman et al., 2015; Kitayama et al., 2011). However, these interactions do not explain how microglia are able to monitor and influence the activity of developing neurons without assessing the overall electrical and metabolic status of the perisomatic neuronal compartment. It is conceivable to hypothesize that microglial interactions with the nucleus-containing cell body would be specifically important for postmitotic neuronal differentiation, cell-fate decisions about neuronal survival or elimination by programmed cell death, as well as the phagocytosis of damaged neuronal cell bodies, all shown to require functional microglia. Furthermore, microglial regulation of the activity of immature neurons devoid of complex synaptic inputs can be essential for proper network formation.

Recently, we discovered specialized morpho-functional interaction sites in the adult brain, named somatic purinergic junctions that are key mediators of microglia-neuron crosstalk (Cserép et al., 2020). These junctions possess a unique ultrastructural and molecular composition that is perfectly suited for bi-directional communication, enabling microglia to readily monitor neuronal status and dynamically influence neuronal functions in the adult brain. In the present study, we tested the hypothesis that somatic purinergic junctions also exist in the developing brain on postmitotic, immature neurons. Moreover, we aimed to determine their precise developmental time course together with the characterization of their nanoscale molecular architecture and subcellular anatomical organization in the context of embryonic, early postnatal and adult neurogenesis.

## Results

To assess microglia-neuron somatic junctions throughout developmental neurogenesis, we used samples from embryonic E15 (ventricular zone/subventricular zone - VZ/SVZ) and early postnatal P1, P8, P15 mice (neocortex). To examine somatic purinergic junctions in association with adult neurogenesis, we analyzed the subgranular layer of the dentate gyrus, one of the two neurogenic niches that persist into adulthood, in P90 mice (Ming and Song, 2011). To discriminate parenchymal microglia from macrophages, microglia were identified based on the expression of the P2Y12 receptor (P2Y12R) and IBA1, which are present in yolk sac-derived microglia from early embryonic days (Hirasawa et al., 2005; Konishi et al., 2017; Mildner et al., 2017), showing an almost perfect colocalization (Figure S1). To visualize immature postmitotic neurons, we performed labeling against doublecortin (DCX, Bennett et al., 2016; Gleeson et al., 1999; Grassivaro et al., 2020; Yoo et al., 2011) (Figure 1.). First, we tested the possible presence of somatic junctions at the two endpoints of the examined timeline. At E15, we observed that microglia were mostly confined to the VZ and SVZ, and rarely present in the cortical plate (Figure 1. A).

**Figure 1.**
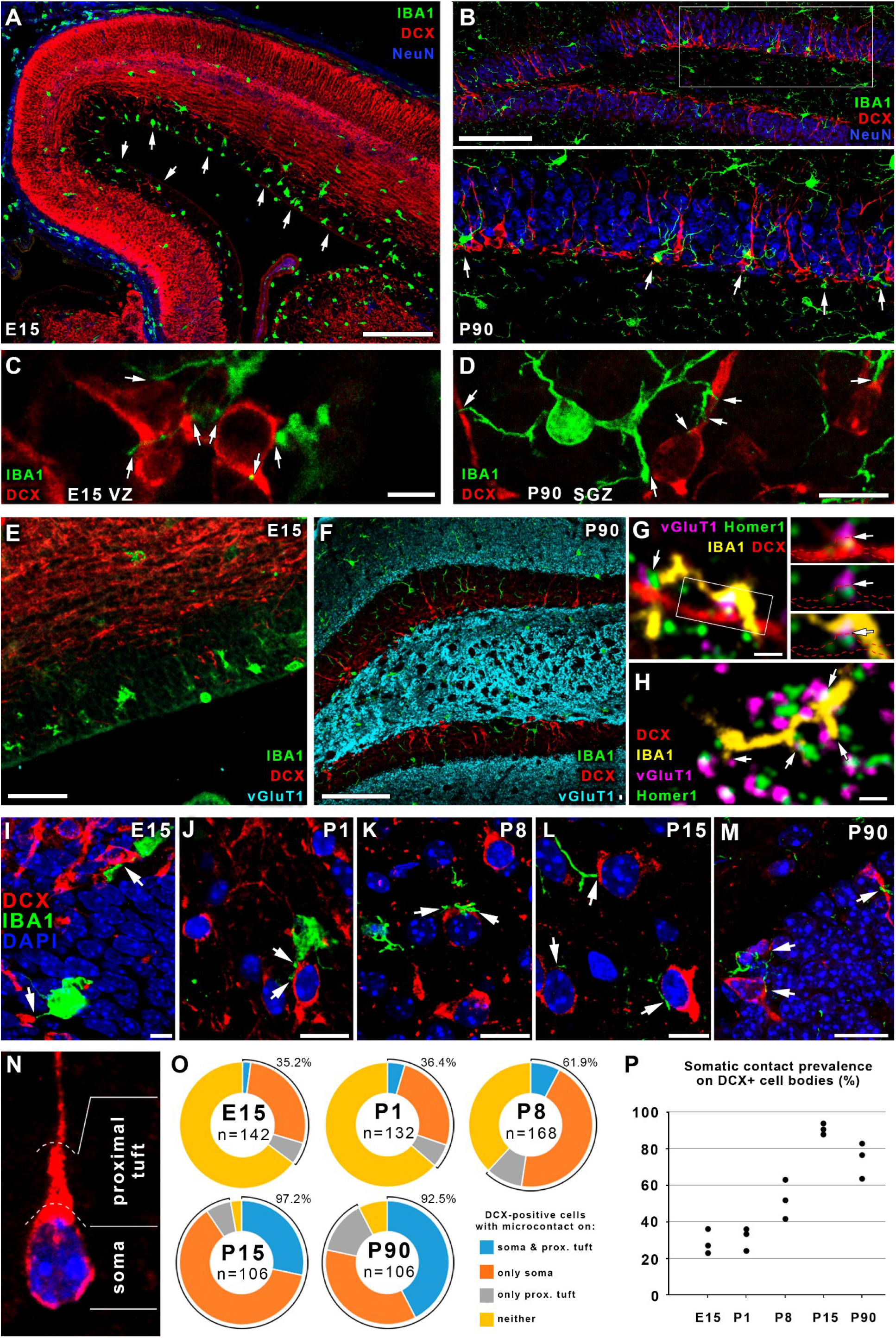
Microglial processes contact the cell bodies, neurites and synapses of DCX-positive postmitotic immature neurons. **A)** Confocal laser scanning microscopic (CLSM) image shows the distribution of IBA1+ microglia (green) in E15 pallium. Postmitotic neurons are labeled for Doublecortin (DCX, red), NeuN is blue, white arrows point to the enrichment of microglia in the SVZ and VZ. **B)** IBA1+ microglia are enriched in the subgranular zone of the DG in P90 mice. Stainings are as on **A**, white arrows point to some microglia. Area within the white box in the upper panel is enlarged in the lower panel. **C-D)** High resolution CLSM images show some examples of microglia-contacts in E15 VZ **(C)** and the DG in P90 mice **(D)**, white arrows point to contact sites. **E)** CLSM image shows the lack of synapses in E15 cortical plate. Samples are labeled for IBA1 (green), DCX (red) and vGluT1 (cyan). **F)** Stainings are the same as on **E**, image shows the abundant presence of glutamatergic synapses outside the granule cell layer. **G)** IBA1 positive microglial processes (yellow) contact developing glutamatergic synapses as identified by the appositions of vGluT1 (magenta) and Homer1 (green) on a DCX+ dendrite (red) in the stratum moleculare of a P90 DG. White arrows point to contacts. **H)** Stainings are same as on **G**, microglial processes contact synapses of mature neurons in the stratum moleculare of a P90 DG. **I-M)** Confocal laser scanning microscopic images show some examples of microglia-neuron somatic contact sites during both developmental and adult neurogenesis. Nuclei are visualized by DAPI (blue), microglia are stained for IBA1 (green) and postmitotic neurons for Doublecortin (DCX, red), white arrows point to contact sites. I – E15, J – P1, K – P8, L – P15, M – P90. **N)** Division of a DCX-positive neuron’s cell body to „soma” and „proximal tuft” compartments. **O)** Ratio of DCX-positive neurons contacted by microglial processes on their somata, proximal tufts or both. Percentage values beside the diagrams represent the ratio of cells that were contacted either way. **P)** Somatic contact prevalence on DCX+ cell bodies (individual values represent different animals). Images and measurements are from the cortical plate in E15 mice, neocortex from P1-P15 mice and hippocampal dentate gyrus from P90 mice, n = 3 mice in each age-group. Scale bar is 200 μm on A, 100 μm on B and F, 5 μm on C, 10 μm on D, J and L, 50 μm on E, 1 μm on G and H, and 15 μm on I, K and M.

In adult mice, we could also detect a strong enrichment of microglia in the subgranular zone of the hippocampal dentate gyrus (Figure 1. B). Importantly, microglial processes frequently contacted both the cell bodies and neurites of DCX+ neurons both during developmental (Figure 1. C) and adult neurogenesis (Figure 1. D). To test whether microglia-synapse contacts could be responsible for microglial process accumulation in the vicinity of developing neuronal cell bodies, we labeled for IBA1, DCX and vGluT1, and confirmed that there are no detectable glutamatergic synapses at E15 (Figure 1. E), and the DG granule cell layer is also devoid of vGluT1+ synapses (Figure 1. F). Nevertheless, co-labeling for IBA1, DCX, vGluT1 and Homer1 revealed that microglial processes contact synapses of both immature (DCX+, Figure 1. G) and mature neurons (Figure 1. H) in the molecular layer of the adult DG. To quantitatively assess the prevalence of somatic junctions, we examined the presence of contacts between microglial processes and 3D reconstructed DCX+ cell bodies through multiple developmental stages (Figure 1. I-P). DCX-positive cells usually possess a main, thick dendritic branch continuous with the cell body, the first short segment (maximum as long as the cell body itself, Figure 1. N), which we termed „proximal tuft”. Somatic contacts were found already at the earliest time point examined (E15), when glutamatergic synapses are absent (Figure 1. E). Importantly, somatic contact prevalence showed a progressive increase during development from more than one-third of developing neurons being contacted at a given time at E15 and P1 to about two-thirds at P8, while at P15 virtually all DCX-positive cells received somatic microglia input. This was also the case during adult neurogenesis in the dentate gyrus of P90 mice (Figure 1. O; 32.5% at E15, 36.4% at P1, 61.9% at P8, 97.2% at P15 and 92.5% at P90; for detailed results see Supplementary Materials/Supplementary Results).

To determine whether the large number of putative direct contacts between microglial processes and DCX-positive immature neurons at the light microscopic level indeed represent somatic purinergic junctions (Cserép et al., 2020) – we first exploited correlated light and electron microscopy (CLEM, Figure 2. A-K).

**Figure 2.**
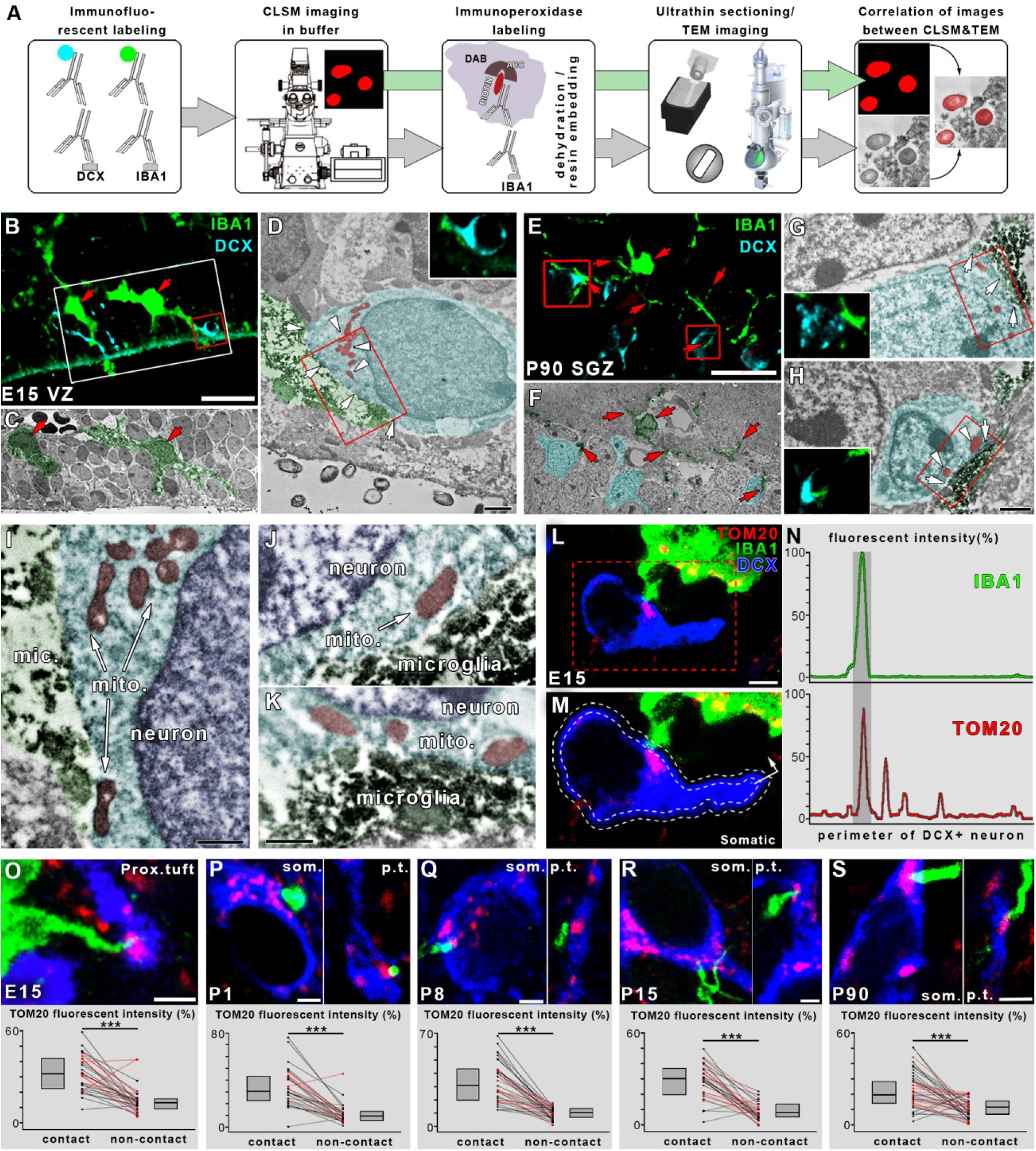
Microglial processes form direct membrane-membrane contacts with the cell bodies of DCX-positive postmitotic immature neurons at sites enriched with mitochondria. **A)** Schematic depiction of correlated light- and electron-microscopy workflow. Immunofluorescent labeling of IBA1 and DCX was followed by CLSM imaging in buffer, after which sections were processed further for immunoperoxidase labeling, dehydration/resin embedding, ultrathin cutting and transmission electron microscopic (TEM) imaging. Finally, the corresponding images from the two imaging modalities were correlated. **B-D)** Maximal intensity projection of a 1.5 μm thick volume from a confocal laser scanning microscopic stack from an E15 mouse shows an example of identified microglia-neuron somatic junctions (**B**). IBA1+ microglia are shown in green, DCX+ neurons are shown in cyan, area in the white box is shown on a correlated TEM image below on **C**, red arrows point to corresponding microglia. Somatic junction within the red box on (**B**) is enlarged on the TEM image on (**D**). White arrows point to the direct membrane-membrane contact, white arrowheads mark neuronal mitochondria close to the junction. Small CLSM insert shows the single confocal image plane closest to the TEM image. TEM images are pseudo-colored (microglia in green, developing neurons in cyan, mitochondria in red). All 6 CLSM-identified contacts proved to be direct membrane-membrane contacts after TEM assessment. **E-H)** Same as (**B-D**) from a P90 mouse, the somatic junction within the left red box on (**E**) is enlarged on the TEM image on (**G**), while the junction within the right red box on (**E**) is enlarged on the TEM image on (**H**). All 10 CLSM-identified contacts proved to be direct membrane-membrane contacts after TEM assessment. **I-K)** Areas within red boxes in D, G and H, respectively, are enlarged from subsequent ultrathin sections. **L**) CLSM image shows an example of an IBA1 labeled (green) microglia contacting the cell body of a DCX-positive postmitotic neuron (blue) exactly where a large mitochondrion (TOM20-labeling, red) resides within the neuronal cell body. Area within the red dashed line is enlarged onto panel **M**. **M**) The process of a semi-automated unbiased analysis of fluorescent intensity area is depicted. White dashed lines represent the outer and the inner profiles, based on the outline of the cell. IBA1 intensity is measured along the outer, TOM20 intensity along the inner profile, starting from the arrow. **N)** The intensity values are plotted along the perimeter of the neuron. Contact site (marked by the darker grey column in the plots) is defined automatically, based on IBA1 fluorescent intensity. **O**) Example of a microglial process (green) contacting the proximal tuft of a DCX-positive neuron in E15 brain. Results show that TOM20 fluorescent intensity is significantly higher within the contact sites than outside them, proving the enrichment of neuronal mitochondria within the microglia-neuron junctions. Each line represents results from one neuron, somatic contacts are represented by black, proximal tuft contacts by red lines. Median values and interquartile range are marked by grey boxes. **P-S**) CLSM examples of mitochondrial enrichment at somatic (som.) and proximal tuft (p.t.) junctions in cortical brain samples from P1, P8, P15 and P90 mice, and corresponding results. Images and measurements are from the cortical plate in E15 mice, neocortex from P1-P15 mice and hippocampal dentate gyrus from P90 mice. Scale bar 30 μm on **B**, 1 μm on **D**, **F-H,**500 nm on **I-K,**20 μm on **E**, 3 μm on **L**, 2 μm on **O-S**.

This approach is ideal to provide direct and diffraction-unlimited evidence that the putative contact sites visualized by the diffraction-limited confocal microscopy are real junctions with nanoscale proximity of the plasma membranes of microglia and neurons. To this end, we performed a combined labeling, and stained DCX and IBA1 with fluorescent immunohistochemistry, while labeling IBA1 with biotinylated antibodies as well (Figure 2. A). By using confocal imaging, we identified putative microglia-neuron contacts (Figure 2. B, E). Subsequently, we developed the immunoperoxidase signal, and processed the samples for electron microscopy. In ultrathin sections, we could then investigate the very same contacts by exploiting diffraction-unlimited resolution of transmission electron microscopy (Figure 2. C-D, F-K).

We reconstructed 6 microglia-neuron contacts from E15 VZ or SVZ, and 10 contacts from P90 dentate gyrus from serial sections. In all cases, we could verify the presence of direct membrane-membrane contacts between the processes and the cell bodies. This CLEM approach also revealed that neuronal mitochondria are frequently enriched at these intercellular contacts (Figure 2. D, G-K), which is also a key morphological feature of somatic purinergic junctions (Cserép et al., 2020). To test whether mitochondrial accumulation at somatic contacts would also be a general feature of DCX-positive cells – as suggested by our electron microscopic results – we applied multiple immunolabeling, CLSM and a semi-automated unbiased method to address this question (Figure 2. L-S). We sampled 3 animals in each age group (n = 164 cells altogether), and found that at each investigated age, TOM20 (mitochondrial protein) immunofluorescent intensity in the cell bodies and proximal tufts of DCX-positive neurons was substantially higher at sites of microglial contact compared with adjacent areas, confirming the strong accumulation of neuronal mitochondria at the interaction sites (for details see Supplementary Materials/Supplementary Results). These results confirmed that neuronal mitochondria are strongly enriched within somatic contact sites established between microglial processes and DCX-positive neuronal cell bodies.

Next, we tested the possibility that enrichment of P2Y12Rs, characteristic microglial signaling molecules is a defining feature of developmental somatic microglia-neuron contacts, as demonstrated earlier in the adult brain (Cserép et al., 2019). We performed multiple immunofluorescent labeling for DCX, IBA1 and P2Y12R and we applied correlated confocal and super-resolution imaging based on Stochastic Optical Reconstruction Microscopy (STORM) (Figure 3. A-F). P2Y12R-immunofluorescence labeling was completely absent in brain sections obtained from two P2Y12R-knockout mice, confirming the specificity of the antibody against P2Y12R (Supp. Figure 2). Next, we performed a quantitative analysis to determine the time course of the somatic purinergic junctions in immature neurons obtained from all five age-groups. We found that virtually all IBA1-positive microglial processes that were in contact with DCX-positive neuronal cell bodies also expressed P2Y12Rs (n = 72 in the VZ and SVZ of E15 developing cortices, n = 60 in the neocortex of P1; n = 85 in P8; n = 74 in P15, and n = 93 in the dentate gyrus of P90 animals; n = 2 mice in each age group, altogether n = 10 mice). Most importantly, correlated confocal and STORM super-resolution imaging confirmed that receptor expression was concentrated strongly on the contacting side of the processes at all age groups examined (n = 35 processes from 5 mice; Figure 3. G; Supplementary Figure 2). This contact-specific clustering of P2Y12 receptors was also a key defining feature of previously described somatic purinergic junctions (Cserép et al., 2020), where they play an important role in intercellular communication.

**Figure 3.**
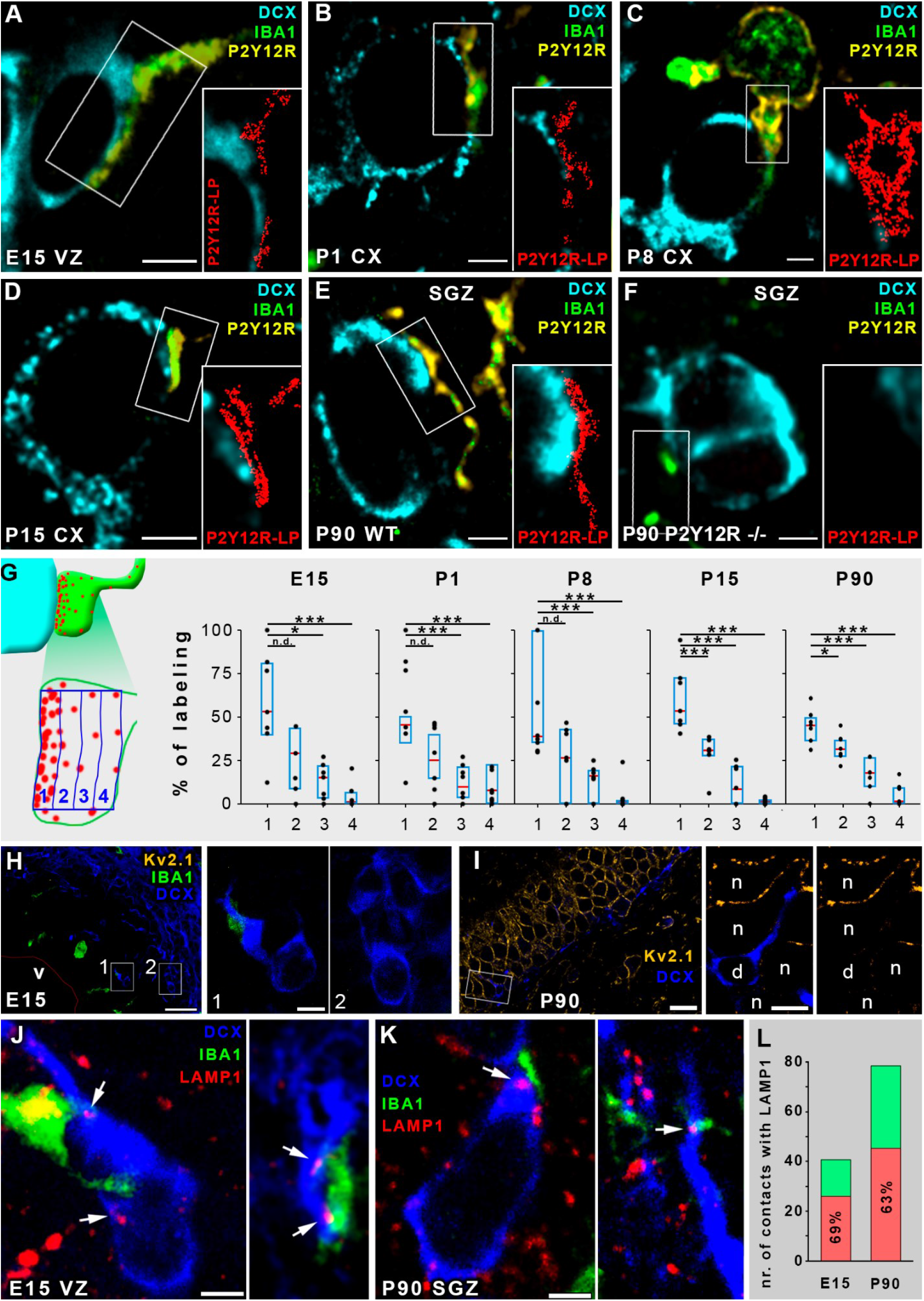
Microglial P2Y12 receptors define somatic purinergic junctions on DCX+ neuronal precursors/immature neurons, while these cells do not express Kv2.1, but contain LAMP1-positve lysosomes in the vicinity of microglial contacts. **A-F)** Correlated STORM super-resolution and CLSM images show the abundant expression of the microglia-specific P2Y12 receptors on microglial processes contacting DCX+ cell bodies. DCX is shown in cyan, IBA1 in green, P2Y12R in yellow, and STORM localization points (LP) for P2Y12R in red on the inserts. Images are from E15 (**A**), P1 (**B**), P8 (**C**), P15 (**D**), P90 (**E**) and P90 P2Y12R-knockout (**F**) mice. Note the complete absence of both the CLSM and the STORM signals for P2Y12R in the knockout mice on (**F**). All IBA1-positive microglial processes in contact with DCX-positive neuronal cell bodies were expressing P2Y12Rs. In the VZ and SVZ of E15 brains n = 72, in the neocortex of P1 n = 60, P8 n = 85, P15 n = 74, and in the dentate gyrus of P90 animals n = 93 contacts were tested (n = 2 mice in each age group, altogether n = 10 mice). **G**) Spatial analysis of super-resolution data shows the enrichment of P2Y12R-labeling on the contacting-side of microglial processes, black dots represent data points, blue bracket is interquartile range, median is shown by red segment; n = 35 processes from 5 mice **H)** CLSM image shows the complete lack of Kv2.1 (yellow) expression in E15 cortical plate (142 fully reconstructed DCX+ cells tested from two mice, zero Kv2.1 positive). DCX (blue), IBA1 (green), ventricle (v) is delineated with thin red line. Rectangular areas labeled with numbers are enlarged on the right. **I)** CLSM image shows a robust expression of Kv2.1 (yellow) in the dentate granule cells of P90 mice, however, DCX-positive (blue) cells are completely void of Kv2.1 labeling (136 fully reconstructed DCX+ cells tested from two mice, zero Kv2.1 positive). Rectangular area is enlarged on the right. Neurons – n, DCX-positive cell – d. **J-K)** CLSM images show examples of IBA1 labeled (green) microglial processes contacting the cell body of a DCX-positive postmitotic neuron (blue) with LAMP1 positive puncta (red, white arrows) in the vicinity. The images are from the SVZ/VZ of E15 (J) and the DG of P90 mice (K). **L)**69% of all contacts in E15 and 63% in P90 contained LAMP1-positive vesicles within the DCX-positive cell bodies in the close vicinity of microglial process contact. Scale bar is 2 μm on A-F, 30 μm on H (5 μm on insets), 15 μm on I (5 μm on insets), 2 μm on J, 2 μm on K.

Our previous studies demonstrated that somatic purinergic junctions are preferentially formed at sites of neuronal exocytosis-promoting Kv2.1 clusters in the adult brain (Cserép et al., 2020). To determine whether the accumulation of this potassium channel is also a defining feature of somatic purinergic junctions during embryonic development, we performed multiple immunofluorescent labeling for DCX and Kv2.1 on samples from E15 and P90 mice (Figure 3. H-I). Notably, we found that Kv2.1 is not expressed in VZ/SVZ at E15 (n = 142 3D reconstructed cells from 2 mice, Figure 3. H). Moreover, although we found a robust expression of Kv2.1 in the mature granule cells in the dentate gyrus in P90 mice, but none of the DCX+ cells expressed Kv2.1 proteins (Figure 3. I; 136 reconstructed cells from two mice), suggesting that alternative routes also exist through which microglia recognize sites of intense exocytosis at the neuronal cell body.

Mitochondria-derived vesicles (MDVs) or other mitochondria-derived cargo often integrate into the endolysosomal pathway (Sugiura et al., 2014), and these vesicles are positive for the lysosomal marker LAMP1 (Soubannier et al., 2012). LAMP1-positive vesicles were identified to be part of mature somatic junctions (Cserép et al., 2020), and we could also observe these organelles in DCX+ cell bodies in the vicinity (within 2 μm) of somatic contacts in the VZ/SVZ of E15 mice (Figure 3. J), and in the DG of P90 mice (Figure 3. K). Quantitative analysis revealed that 69% and 63% of the examined somatic contacts contained adjacent LAMP1+ puncta in E15 (n = 49 contacts from 2 mice), and in P90 (n = 71 contacts from 2 mice, Figure 3. L) mice, respectively, suggesting that cellular content could also be released at developmental somatic junctions from immature neurons.

## Discussion

In the present study, we provide direct molecular and anatomical evidence that specialized contact sites are present between microglia and immature postmitotic neurons throughout embryonic and postnatal development and even during adult neurogenesis. These contact sites have ideal nanoscale architecture to underlie microglial regulation of neuronal development. Direct microglia-neuron interactions have already been implicated in various developmental processes. However, these synaptic/dendritic and axonal interactions could not fully explain the entire spectrum of the diverse developmental roles of microglia, as the majority of cell-fate decisions are linked to cell bodies (Donato et al., 2019; Hobert et al., 2010; Terenzio et al., 2017), necessitating the interactions between microglial processes and the somatic compartment. The relevance of communication between microglia and the somatic compartment of developing neurons is even more prominent, knowing that microglial regulation of neuronal development at early stages takes place in cells that are yet devoid of synapses. Indeed, we showed that direct somatic microglia-neuron contacts persist in all stages of rodent development as well as during adult neurogenesis, while the ratio of contacted DCX+ cell bodies showed a substantial increase during postnatal development, with a sharp rise between P1 and P8, matching the most intensive period of synaptogenesis (Fiala et al., 1998; Khazipov et al., 2001; Tyzio et al., 1999), the switching of depolarizing GABA effects to hyperpolarizing, and robust changes in network activity patterns (Ben-Ari et al., 2012; Kaila et al., 2014). We suggest that through these contacts microglia can readily monitor the status and function of neurons as demonstrated in the adult brain, allowing microglia to dynamically influence neuronal functions and cell-fate decisions through robust bi-directional communication (Cserép et al., 2020), and assist activity-dependent neuronal integration.

The role of neuronal mitochondria in developmental and adult neurogenesis has become evident in the last years (Arrázola et al., 2019; Khacho and Slack, 2018; Rangaraju et al., 2019). Mitochondrial metabolism fundamentally regulates developmental and adult neurogenesis and differentiation (Beckervordersandforth, 2017; Lorenz and Prigione, 2017), while mitochondrial structural dynamics of postmitotic neurons has been shown to drive developmental neurogenesis (Iwata et al., 2020), and to be instrumental regarding cell-fate decisions at the same time (Bhola and Letai, 2016). Mitochondria are also critical for injury-related responses in the developing brain (Hagberg et al., 2014), through the induction of programmed cell death (Yamaguchi and Miura, 2015) or the morphogenesis of neurons (Kimura and Murakami, 2014). Thus, somatic mitochondria may provide ultimate readout for microglia and function as possible sites to influence neuronal physiology or cell death as demonstrated in the adult brain (Chandel, 2014; Whelan and Zuckerbraun, 2013). We found a strong association between somatic mitochondria of DCX+ cells and the contact sites formed by microglial processes during both development and adult neurogenesis, similar to that seen in mature neurons (Cserép et al., 2020). Microglia are thus in an ideal position at somatic junctions to sense mitochondria-derived signals – including purine metabolites (Bajwa et al., 2019; Cserép et al., 2020; Davalos et al., 2005; Gouveia et al., 2017) – and influence cellular functions. Through these junctions, microglia could modulate the activity or morphogenesis of postmitotic cells via altering their metabolism or mitochondrial function.

Mitochondria-derived vesicles, other mitochondria-related substances or signaling molecules released from neuronal cell bodies are carrying important information of the source-cell’s overall status. The somato-dendritic region of developing neurons has been shown to be responsible for the majority of the exocytotic events (Urbina and Gupton, 2020; Urbina et al., 2018). In the case of mature purinergic junctions, Kv2.1-channel clusters seem to be important regulators of somatic exocytosis (Mohapatra et al., 2007). Although we could not detect the expression of these Kv2.1 clusters in DCX-positive cells, there could be other delayed rectifier KV-proteins present, or several other molecular scaffolds providing sufficient exocytotic surface, like it is suggested in the case of mature GABAergic neurons, which do not express Kv2.1 channels, but still have somatic junctions formed by microglial processes. Our results, showing the presence of LAMP1-positive lysosomes in the close vicinity of somatic junctions on DCX-positive cell bodies suggest that cellular content could be released here, acting as a local readout for microglial processes, which are assessing cellular status. Furthermore, vesicular nucleotide transporter, which is responsible for packing purine nucleotides into vesicles and has been functionally linked to somatic neuronal ATP release, has also been shown to be expressed in DCX-positive developing neurons, colocalizing with LAMP1-positive vesicles (Cserép et al., 2020; Menéndez-Méndez et al., 2017). This suggests that among the wide variety of signaling molecules and mitochondria-related substances, which are presumably released at these somatic junctions, purinergic nucleotides are key candidates.

Purinergic signaling and microglial P2Y12 receptors have been implicated in the regulation of neurogenesis and brain development (Ribeiro et al., 2019; Rodrigues et al., 2019). We found a robust expression of P2Y12R-s on microglial processes involved in the somatic junctions throughout development and during adult neurogenesis. These microglial receptors are also expressed during human development (Mildner et al., 2017), suggesting that this feature is evolutionary conserved. Microglial P2Y12 signaling has been shown to regulate neurogenesis and immature neuronal projections (Mo et al., 2019), and to promote proliferation of adult mouse SVZ cells (Suyama et al., 2012). On the other hand, microglia couple phagocytosis with the progression of apoptosis via P2Y12R signaling during development (Blume et al., 2020). All these data suggest a role for purinergic and P2Y12R-dependent signaling processes during the microglial regulation of neurodevelopment. Since these receptors are robustly expressed on microglial processes in the somatic junctions, and these contact sites have been shown to release purinergic metabolites in mature neurons (Cserép et al., 2020), it is likely that similar signaling pathways within these junctions could be involved in the microglial regulation of neurodevelopment. In mature somatic junctions, microglial processes sense activity-dependent somatic ATP-release (Cserép et al., 2020). During the developmental program of neurons, basic changes in activity patterns parallel the most active period of synaptogenesis (Fiala et al., 1998). During the first postnatal week, depolarizing GABAergic effects switch to hyperpolarizing and glutamatergic synapses appear, while the primitive synchronous developmental oscillatory activity gradually changes to more mature activity patterns (Ben-Ari et al., 2012). Monitoring and regulation of neuronal activity by microglia through somatic junctions may provide means to exert developmental regulation during this critical period, and assist the formation of complex neuronal networks. This idea is also supported by the steep increase of somatic contact prevalence exactly during the first postnatal week.

Our findings raise several questions and require several functional confirmation studies, which are beyond the scope of the present paper. First of all, we focused our experiments on postmitotic immature neurons, confirmed by their DCX expression. However, microglial regulation of neurogenesis, proliferation, differentiation and survival of neural precursors or migration are active processes even at embryonal stages (Aarum et al., 2003b; Bilimoria and Stevens, 2015; Cunningham et al., 2013; Erblich et al., 2011; Ueno et al., 2013), often affecting more immature progenitors. Nevertheless, somatic interactions have already been observed between microglial processes and the cell bodies of dividing neuronal progenitors (Cunningham et al., 2013; Noctor et al., 2019; Penna et al., 2020), thus, it needs to be addressed whether these are also functioning as somatic junctions. Secondly, at this stage we concentrated on the molecular and structural assessment of somatic microglia-neuron junctions, since to extend our experiments towards functional directions we would either need to manipulate immature neurons at different maturational stages, or to apply microglia-specific interventions at multiple developmental stages, clearly surpassing the scope of this study. Longitudinal *in vivo* imaging studies are also necessary to decipher whether developmental phagocytosis (Diaz-Aparicio et al., 2020; Sierra et al., 2010; VanRyzin et al., 2019) would require somatic junctions as specific checkpoints. Also, microglia-specific interventions should be used to explore, whether the differentiation, migration or integration of progenitors / developing neurons is regulated through somatic junctions. Mitochondrial changes are involved in developmental apoptosis of neuronal progenitors (László et al., 2020), and these changes could also be sensed or even regulated by microglia, which can in turn eliminate unfit progenitors via inducing programmed cell death followed by phagocytosis. The NAD+:NADH ratio plays an important role in regulating stem cell-fate (Khacho and Slack, 2018), and we showed previously that microglia are able to regulate mitochondrial NADH-production at somatic junctions, in a P2Y12R-dependent manner (Cserép et al., 2020). This also raises the possibility that microglia could – at least in part – exert developmental regulation through modifying the mitochondrial function of neuronal progenitors or immature neurons.

Microglia populate the brain early during development (Ginhoux et al., 2010; Verney et al., 2010) and maintain a self-renewing, long-lived population thereafter (Füger et al., 2017; Réu et al., 2017) with marked transcriptome changes throughout life (Hammond et al., 2019; Matcovitch-Natan et al., 2016). How micorglia shape neurodevelopment under different conditions via different interactions will need to be investigated in the future. Since even transient perturbations of developmental microglia-neuron interactions can lead to complex and persisting disruptions in neuronal networks (Paolicelli and Ferretti, 2017; Park et al., 2020; Prins et al., 2018; Xu et al., 2020), the clinical relevance – especially in neurodevelopmental disorders – to achieve a better understanding of the underlying complex neuro-immune communication and compartment-specific microglia-neuron interactions in health and disease is evident.

## Methods

### Animals

Experiments were carried out on E15, P1, P8, P15 and P90 old C57BL/6 (RRID:IMSR_JAX:000664) mice. Animals were bred at the SPF unit of the Animal Care Unit of the Institute of Experimental Medicine (IEM, Budapest, Hungary). Mice had free access to food and water and were housed under light-, humidity- and temperature-controlled conditions. All experiments were performed in accordance with the Institutional Ethical Codex and the Hungarian Act of Animal Care and Experimentation guidelines (40/2013, II.14), which are in concert with the European Communities Council Directive of September 22, 2010 (2010/63/EU). The Animal Care and Experimentation Committee of the Institute of Experimental Medicine and the Animal Health and Food Control Station, Budapest, have also approved the experiments under the number PE/EA/1021-7/2019, PE/EA/673-7/2019. All experiments were performed in accordance with ARRIVE guidelines.

### Perfusion and tissue processing for histology

Adult mice were anesthetized by intraperitoneal injection of 0.15-0.25 ml of an anaesthetic mixture (containing 20 mg/ml ketamine, 4 mg/ml xylazine-hydrochloride). Animals were perfused transcardially with 0.9% NaCl solution for 1 minute, followed by 4% freshly depolymerized paraformaldehyde (PFA) in 0.1 M phosphate buffer (PB) pH 7.4 for 40 minutes, and finally with 0.1 M PB for 10 minutes to wash the fixative out. Blocks containing the primary somatosensory cortex and dorsal hippocampi were dissected and coronal sections were prepared on a vibratome (VT1200S, Leica, Germany) at 50 μm thickness for immunofluorescent experiments and electron microscopy. For some experiments, 25 μm thick sections from E15 mouse brains were prepared on a cryostat, and dried onto glass slides.

### Immunofluorescent labeling and confocal laser scanning microscopy (CLSM)

Before the immunofluorescent staining, the 50 μm thick sections were washed in PB and Tris-buffered saline (TBS). This was followed by blocking for 1 hour in 1% human serum albumin (HSA; Sigma-Aldrich) and 0.03-0.1% Triton X-100 dissolved in TBS and 200 ug/ml Digitonin (D141-100MG, Sigma). After this, sections were incubated in mixtures of primary antibodies overnight at room temperature. After incubation, sections were washed in TBS and were incubated overnight at 4 °C in the mixture of secondary antibodies, all diluted in TBS. Secondary antibody incubation was followed by washes in TBS, PB, the sections were mounted on glass slides, and coverslipped with Aqua-Poly/Mount (Polysciences). Immunofluorescence was analyzed using a Nikon Eclipse Ti-E inverted microscope (Nikon Instruments Europe B.V., Amsterdam, The Netherlands), with a CFI Plan Apochromat VC 60X oil immersion objective (numerical aperture: 1.4) and an A1R laser confocal system. We used 405, 488, 561 and 647 nm lasers (CVI Melles Griot), and scanning was done in line serial mode, pixel size was 50×50 nm. Image stacks were obtained with NIS-Elements AR. For primary and secondary antibodies used in this study, please see Table 1.

**Table 1.** List of antibodies used in the study.

| **Primary antibodies** | **host** | **source** | **catalog nr.** | **RRID** |
| --- | --- | --- | --- | --- |
| DCX | chicken | Synaptic Systems | 326-006 | RRID: AB_2737040 |
| IBA1 | guinea pig | Synaptic Systems | 234004 | RRID: AB_2493179 |
| IBA1 | goat | Novusbio | NB100- | RRID:AB_521594 |
| IBA1 | rabbit | Wako Chemicals | 019-19741 | RRID: AB_839504 |
| Kv2.1 | mouse | NeuroMab | 75-014 | RRID:AB_10673392 |
| LAMP1 | rabbit | Abcam | ab24170 | RRID:AB_775978 |
| P2Y12R | rabbit | Anaspec | AS-55043A | RRID:AB_2298886 |
| TOM20 | rabbit | Santa Cruz | sc-11415 | RRID:AB_2207533 |
| **Secondary antibodies** | **host** | **source** | **catalog nr.** | **RRID** |
| biotinylated anti-rabbit | donkey | BioRad | 644008 | RRID:AB_619842 |
| Alexa 488 anti-goat | donkey | Jackson | 705-546-147 | RRID:AB_2340430 |
| Alexa 488 anti-guinea-pig | donkey | Jackson | 706-546-148 | RRID: AB_2340473 |
| Alexa 594 anti-rabbit | donkey | LifeTech | A21207 | RRID:AB_141637 |
| Alexa 594 anti-chicken | donkey | Jackson | 703-586-155 | RRID: AB_2340378 |
| Alexa 647 anti-guinea-pig | donkey | Jackson | 706-606-148 | RRID:AB_2340477 |
| Alexa 647 anti-chicken | donkey | Jackson | 703-606-155 | RRID: AB_2340380 |
| Alexa 647 anti-rabbit | donkey | Jackson | 711-605-152 | RRID:AB_2492288 |
| Alexa 647 anti-mouse | donkey | Jackson | 715-605-150 | RRID:AB_2340866 |

### Quantitative analysis of CLSM data

Quantitative analysis of each dataset was performed by at least two observers, who were blinded to the origin of the samples, the experiments and did not know of each other’s results. For the colocalisation measurement of microgial markers during development in the mouse brain, confocal stacks with triple immunofluorescent labeling (IBA1-Gp, IBA1-Gt and P2Y12R) were used, acquired from the VZ/SVZ region of E15, neocortex of P1-P15 and dentate gyrus of P90 mice. During the analysis cells with a DAPI-labeled nucleus in the IBA1-Gp channel were randomly selected. After that we measured the colocalisation with the other two microglia labelings (IBA1-Gt, P2Y12R) in each age group.

For the analysis of somatic junction prevalence, confocal stacks with double immunofluorescent labeling (DCX and IBA1) were acquired from the VZ/SVZ region of E15, neocortex of P1-P15 and dentate gyrus of P90 mice. All labeled and identified cells or proximal tufts (the proximal segment of the thickest dendrite of the cell, not longer than the cell body itself) were counted, when the whole cell body or tuft was located within the Z-stack. Given somata or proximal tufts were considered to be contacted by microglia, when a microglial process clearly touched it (i.e. there was no space between neuronal soma and microglial process) on at least 0.5 μm long segment.

TOM20 fluorescent intensity profiles were analyzed using a semi-automatic method. Confocal stacks with triple immunofluorescent labeling (microglia, DCX and TOM20) were collected. The section containing the largest cross-section of a neuron was used to trace the cell membrane according to DCX-labeling. This contour was then expanded and narrowed by 0.5 μm to get an extracellular and an intracellular line, respectively (Figure 4B). The intensity of fluorescent labeling was analyzed along these lines. After normalizing and scaling, microglial contact was identified where microglial fluorescent intensity was over 20% of total, for at least 500 nm. Then the contact area was extended 500-500 nm on both sides, and TOM20 fluorescent intensity within these areas was measured for “contact” value.

### STORM super-resolution imaging

Free-floating brain sections were blocked with 1% human serum albumin followed by immunostaining with rabbit anti-P2Y12R, guinea-pig anti IBA1 and chicken anti-DCX antibodies, followed by anti-rabbit Alexa 647, anti-guinea-pig Alexa 488 and anti-chicken Alexa 594 secondary antibodies. Sections were mounted onto #1.5 borosilicate coverslips and covered with imaging medium containing 5% glucose, 0.1 M mercaptoethylamine, 1 mg/ml glucose oxidase, and catalase (Sigma, 1500 U/ml) in Dulbecco’s PBS (Sigma), immediately before imaging. STORM imaging was performed for P2Y12R (stimulated by a 647 nm laser) by using a Nikon N-STORM C2+ superresolution system that combines ‘Stochastic Optical Reconstruction Microscopy’ technology and Nikon’s Eclipse Ti research inverted microscope to reach a lateral resolution of 20 nm and axial resolution of 50 nm. Imaging was performed using the NIS-Elements AR 4.3 with N-STORM 3.4 software. Molecule lists were exported from NIS in txt format, and the three image planes of the ics-ids file pairs from the deconvolved confocal stacks matching the STORM volume were converted to the ome-tiff format using FIJI software. Confocal and corresponding STORM images were fitted in Adobe Photoshop CS6. Localization points exceeding a photon count of 2000 were counted as specific superresolution localization points. Local density filter (10 neighbours within 150 nm for P2Y12R) and Z-filter (±300 nm from focal plane) was applied to the localization points.

### Correlated CLSM and immune-electron microscopy

After extensive washes in PB and 0.05 M TBS sections were blocked in 1% HSA in TBS and 0,03% Triton X.100 and 50 ug/ml digitonin (D141-100MG, Sigma). Then, they were incubated in primary antibodies (chicken anti DCX, guinea-pig anti IBA1, together with either rabbit anti-IBA1 or rabbit anti-P2Y12R) diluted in TBS containing 0.05% sodium azide for 2-3 days. After repeated washes in TBS, the sections were incubated in fluorophore-conjugated and biotinylated secondary antibodies diluted in TBS. Secondary antibody incubation was followed by washes in TBS and PB. Sections were mounted in PB, coverslipped, sealed with nail-polish. Immunofluorescence for DCX and IBA1-Gp was analysed using a Nikon Eclipse Ti-E inverted microscope (Nikon Instruments Europe B.V., Amsterdam, The Netherlands), with a CFI Plan Apochromat VC 60X water immersion objective (NA: 1.2) and an A1R laser confocal system. We used 488, 561 and 642 nm lasers, and scanning was done in line serial mode. Image stacks were obtained with NIS-Elements AR software with a pixel size of 50×50 nm in X-Y, and 150 nm Z-steps. After imaging, sections were washed thoroughly, and incubated in avidin–biotinylated horseradish peroxidase complex (Elite ABC; 1:300; Vector Laboratories) diluted in TBS for 3 h at room temperature or overnight at 4 °C. The immunoperoxidase reaction against IBA1-Rb or P2Y12R was developed using 3,3-diaminobenzidine (DAB; Sigma-Aldrich) as chromogen. The sections were then treated with 1% OsO_4_ in 0.1 M PB, at room temperature, dehydrated in ascending alcohol series and in acetonitrile and embedded in Durcupan (ACM; Fluka). During dehydration, the sections were treated with 1% uranyl acetate in 70% ethanol for 20 min. For electron microscopic analysis, correlated tissue samples from the VZ of E15 and from the gyrus dentatus of P90 mice were glued onto Durcupan blocks. Consecutive 70 nm thick sections were cut using an ultramicrotome (Leica EM UC7) and picked up on Formvar-coated single-slot grids. Ultrathin sections were examined in a Hitachi 7100 electron microscope equipped with a Veleta CCD camera (Olympus Soft Imaging Solutions, Germany). During the correlation a large number of wildfield images were taken from the samples and several maps were constructed to ensure that the very same tissue volumes are processed for electron microscopy that have been imaged using CLSM. Somatic junctions were randomly selected in the confocal stacks, and these contacts were found on the electron microscopic images to test the prevalence of direct membrane-membrane contacts. For correlation and montage stiching Adobe Photshop CS6 and Fiji were used.

### Statistical analysis

All quantitative assessment was performed in a blinded manner wherever it was possible. Based on the type and distribution of data populations (examined with Shapiro-Wilks W test) we applied the nonparametric Wilcoxon signed rank test to compare two dependent data groups. Statistical analysis was performed with the Statistica 13.4.0.14 package (TIBCO), differences with p<0.05 were considered significant throughout this study.

## Data availability

No omics datasets were generated or analyzed during the current study.

## Acknowledgements

We thank Drs. László Barna, Pál Vági and the Nikon Imaging Center at the Institute of Experimental Medicine for kindly providing microscopy support, and Dóra Gali-Györkei, Balázs Pintér and Erika Tischler for their excellent technical assistance.

## Author contributions

C.C., B.P., Z.I.L., Z.L. and A.D. conceived the project. Tissue samples were prepared by Z.I.L. and Z.L.; immunohistochemistry and confocal microscopy was performed by C.C., B.P., and A.D.S; STORM microscopy was performed by B.P.; correlated light and electron microscopy was performed by C.C. and A.D.S.; analysis was performed by C.C., B.P., A.D.S., A.K. and M.N; I.K., provided resources and essential intellectual contribution, and revised the manuscript. A.D. obtained funding and supervised the project. C.C. and A.D. wrote the paper with input from all authors.

## Supplementary Information

### Supplementary results

**Related to Figure 1. I-O, (contact analysis)**. 35.2% of the DCX+ neurons received microglial process contact in the VZ and SVZ at E15 (2.1% was contacted both on the soma and proximal tuft, 27.5% only on the soma and 5.6% only on the prox. tuft, n = 142 cells from 3 mice; Figure 1I, O). At P1, 36.4% of the DCX+ neurons received microglial process contact in the neocortex (4.5% was contacted both on the soma and proximal tuft, 25.8% only on the soma and 6.1% only on the prox. tuft, n = 132 cells from 3 mice; Figure 1J, O). At P8, 61.9% of the DCX+ neurons received microglial process contact in the neocortex (7.7% was contacted both on the soma and proximal tuft, 44.6% only on the soma and 9.5% only on the prox. tuft, n = 168 cells from 3 mice; Figure 1K, O). At P15, 97.2% of the DCX+ neurons received microglial process contact in the neocortex (28.3% was contacted both on the soma and proximal tuft, 62.2% only on the soma and 6.6% only on the prox. tuft, n = 106 cells from 3 mice; Figure 1L, O). At P90, 92.5% of the DCX+ neurons received microglial process contact in the dentate gyrus (42.5% was contacted both on the soma and proximal tuft, 35.8% only on the soma and 14.2% only on the prox. tuft, n = 106 cells from 3 mice; Figure 1M, O). These data confirm that a significant proportion of postmitotic neurons receive somatic contacts from microglial processes at any given timepoint, and this proportion is growing towards adulthood.

**Related to Figure 2. I-O (TOM20 analysis).** TOM20 fluorescent intensity was 146% higher in contact-areas than in non-contact areas in E15 mice (Figure 2I-K, p<0.001; n = 30 contacts; 0.32, 0.22-0.42 vs. 0.13, 0.09-0.15 median and interquartile range in contact and non-contact areas, respectively), 233% higher in P1 mice (Figure 2L, p<0.001; n = 30 contacts; 0.3, 0.23-0.43 vs. 0.09, 0.05-0.13), 233% higher in P8 (Figure 2M, p<0.001; n = 33 contacts; 0.3, 0.19-0.42 vs. 0.09, 0.06-0.13), 275% higher in P15 mice (Figure 2N, p<0.001; n = 30 contacts; 0.3, 0.21-0.36 vs. 0.08, 0.05-0.12) and 120% higher in P90 mice (Figure 2O, p<0.001; n = 41 contacts; 0.22, 0.15-0.31 vs. 0.1, 0.06-0.15).

**Related to Figure 3L (super-resolution distribution analysis)**. The 1^st^ quarter of the microglial process is touching the DCX-positive cell, the 4^th^ is the opposite side. E15: percentage of labeling, median values (med.) 1^st^: 52, 2^nd^: 29, 3^rd^: 15, 4^th^: 0, p-values: 1 vs. 2: 0.0966, 1 vs. 3: 0.0106, 1 vs. 4: 0.0033; P1: med. 1^st^: 44, 2^nd^: 24, 3^rd^: 9, 4^th^: 7, p: 1 vs. 2: 0.0966, 1 vs. 3: 0.0033, 1 vs. 4: 0.0033; P8: med. 1^st^: 38, 2^nd^: 26, 3^rd^: 16, 4^th^: 0, p: 1 vs. 2: 0.1252, 1 vs. 3: 0.0022, 1 vs. 4: 0.0022; P15: med. 1^st^: 53, 2^nd^: 30, 3^rd^: 8, 4^th^: 0, p: 1 vs. 2: 0.0022, 1 vs. 3: 0.0022, 1 vs. 4: 0.0022; P90: med. 1^st^: 46, 2^nd^: 32, 3^rd^: 18, 4^th^: 1, p: 1 vs. 2: 0.0409, 1 vs. 3: 0.0022, 1 vs. 4: 0.0022; all comparisons are done with the Mann-Whitney U test.

**Supplementary Figure 1.**
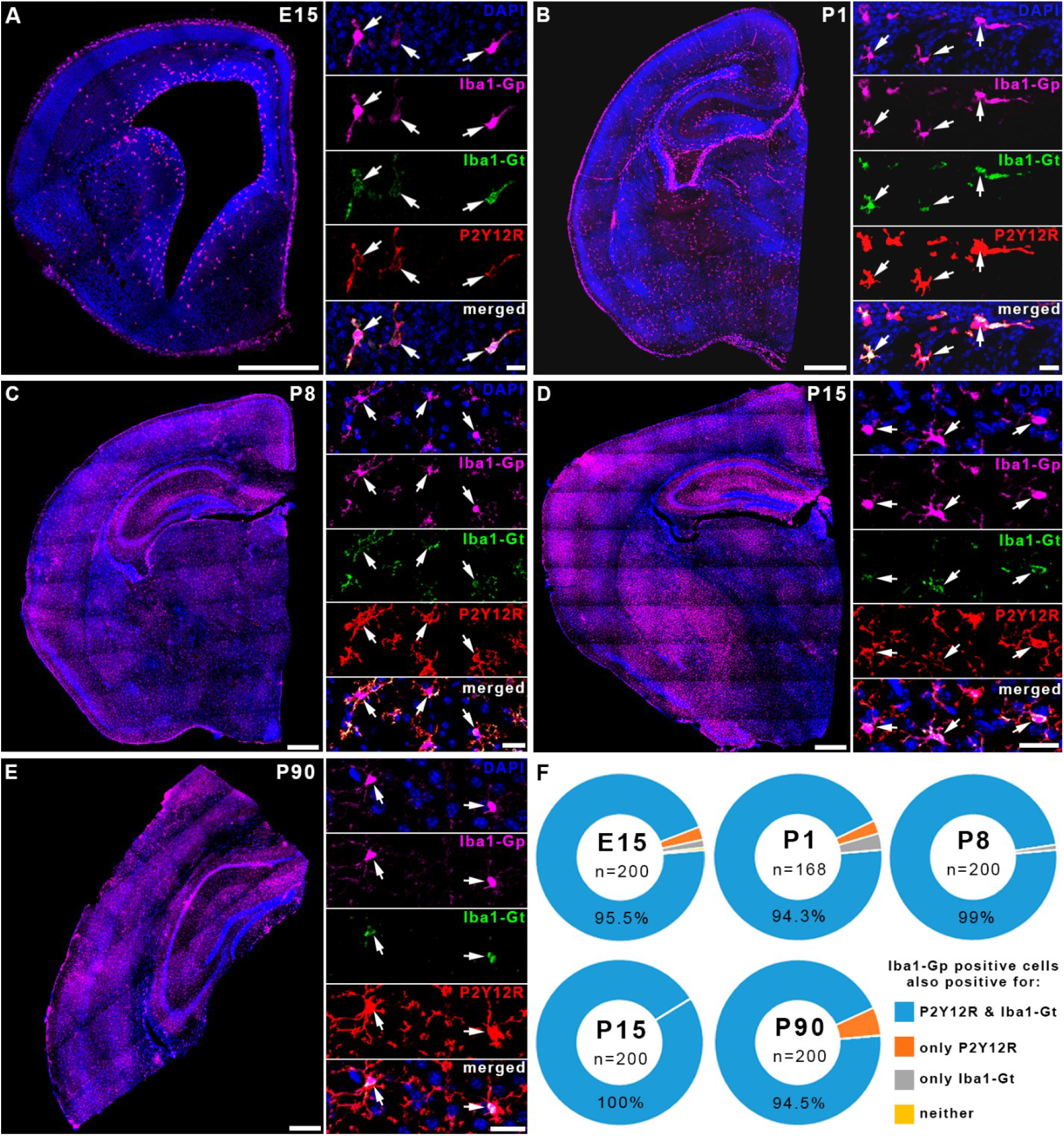
Colocalisation of microgial markers during development in the mose brain. **A-E)** Large montage images obtained with confocal laser scanning microscopy show the ovarall distribution pattern of IBA1-positive cells in mouse brains. Inserts show colocalisation of markers (magenta: IBA1-guinea-pig ab., green: IBA1-goat ab., red: P2Y12R, blue: DAPI), white arrows point to microglial cell bodies. A – E15, B – P12, C – P8, D – P15, E – P90. **F)** The wast majority of IBA1-Gp positive cells were also positive for both IBA1-Gt and P2Y12R. Insert images and measurements are from the cortical plate in E15 mice, neocortex from P1-P15 mice and hippocampal dentate gyrus from P90 mice. Scale bars on large images: 500 μm, on inserts 25 μm.

**Supplementary Figure 2.**
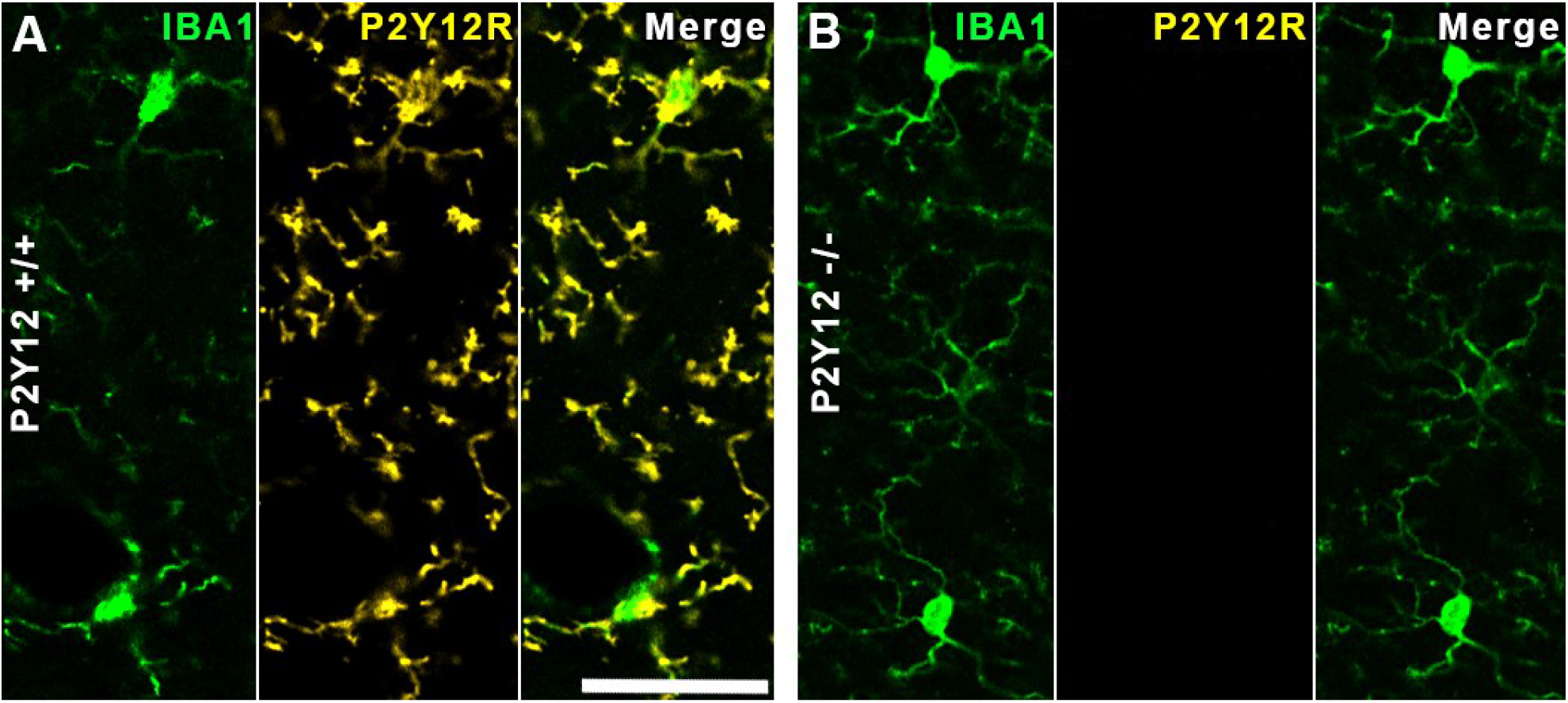
Colocalisation of microgial markers IBA1 and P2Y12R in WT and P2Y12R-KO mice. **A)** Confocal laser scanning microscopy show colocalisation of IBA1 (green) and P2Y12R (yellow) in WT mouse brain. **B)** Same as on A, but on P2Y12R-KO mouse brain. P2Y12R-labeling is completely absent in the KO animals. Scale bar is 10 μm.

